# X-ray-driven chemistry and conformational heterogeneity in atomic resolution crystal structures of bacterial dihydrofolate reductases

**DOI:** 10.1101/2023.11.07.566054

**Authors:** Nathan Smith, Alexander R. Horswill, Mark A. Wilson

## Abstract

Dihydrofolate reductase (DHFR) catalyzes the NADPH-dependent reduction of dihydrofolate to tetrahydrofolate. Bacterial DHFRs are targets of several important antibiotics as well as model enzymes for the role of protein conformational dynamics in enzyme catalysis. We collected 0.93 Å resolution X-ray diffraction data from both *Bacillus subtilis* (Bs) and *E. coli* (Ec) DHFRs bound to folate and NADP^+^. These oxidized ternary complexes should not be able to perform chemistry, however electron density maps suggest hydride transfer is occurring in both enzymes. Comparison of low- and high-dose EcDHFR datasets show that X-rays drive partial production of tetrahydrofolate. Hydride transfer causes the nicotinamide moiety of NADP^+^ to move towards the folate as well as correlated shifts in nearby residues. Higher radiation dose also changes the conformational heterogeneity of Met20 in EcDHFR, supporting a solvent gating role during catalysis. BsDHFR has a different pattern of conformational heterogeneity and an unexpected disulfide bond, illustrating important differences between bacterial DHFRs. This work demonstrates that X-rays can drive hydride transfer similar to the native DHFR reaction and that X-ray photoreduction can be used to interrogate catalytically relevant enzyme dynamics in favorable cases.

## Introduction

Dihydrofolate reductase (DHFR) catalyzes the NADPH-dependent reduction of 7,8-dihydrofolate (DHF) to 5,6,7,8-tetrahydrofolate (THF) (Fig. 1A). DHFR activity is critical for folate-dependent biochemistry, including nucleoside biosynthesis and 1-carbon processes connected with vitamin B12 (cobalamin)[1]. Because of DHFR’s importance for de novo synthesis of nucleotides and certain amino acids in proliferating cells, it is the target of several drugs, including methotrexate and trimethoprim [2,3].

**Figure 1.**
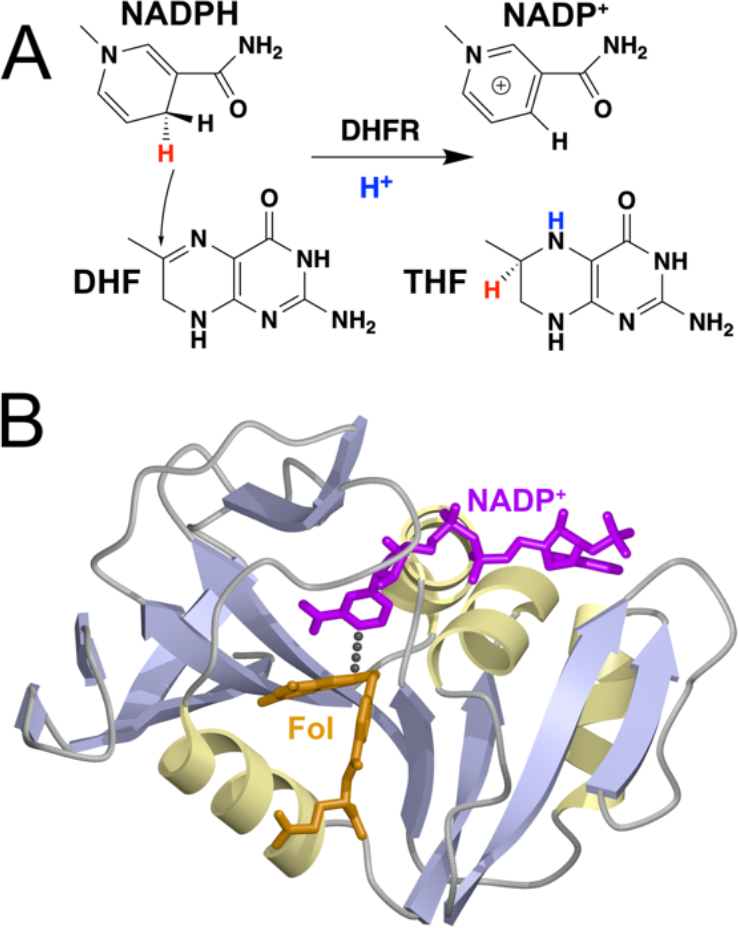
DHFR reaction scheme and structure of the EcDHFR-NADP^+^-folate abortive ternary complex. (A) shows the hydride (red) donation by NADPH to dihydrofolate (DHF) and protonation (blue) to form the product tetrahydrofolate (THF). (B) shows a ribbon diagram of EcDHFR in complex with NADP^+^ (purple) and folate (Fol; orange). The dotted line connects the C4 atom in NADP^+^ to the C6 atom in folate, which represents the hydride transfer pathway in the true NADPH-DHF Michaelis complex.

*E. coli* DHFR (EcDHFR) is the most extensively studied example of this family of enzymes (Fig. 1B), with X-ray crystal structures representing all proposed intermediates along the reaction coordinate having been determined [4]. A complete kinetic scheme for the enzyme was developed in the 1980s [5] and detailed biophysical characterization using NMR [6-9], spectroscopy, [10-12], isotope effects, [13-17], and computation [18-22] followed. Among other results, these studies showed that an active site loop containing Met20 samples multiple states in the apo, cofactor-bound, ternary (Michaelis), and the product-bound EcDHFR complexes. The three principal conformations of the Met20 loop (open, closed, and occluded) gate access of the DHF substrate to the active site, assist release of the THF product and oxidized NADP^+^ cofactor, and couple to larger subdomain motions in EcDHFR [4]. Even in the closed conformation, the dynamics of the Met20 sidechain has been proposed to play an important role in substrate protonation [20,22-24]. Neutron crystal structures of the abortive ternary EcDHFR-folate-NADP^+^ complex at multiple pH values clarified how Met20 sidechain dynamics can provide transient water access to the active site to protonate the substrate [25]. Despite the importance of Met20 motion for sampling catalytically productive conformations of EcDHFR, Met20 is not highly conserved in DHFRs [26]. In human DHFR it is substituted by Leu22 [27] and in *B. anthracis* DHFR it is Leu21 [28]. The poor conservation of methionine at this position suggests that Met20 loop dynamics are not essential for the chemical step itself [29] and that the details of substrate protonation during catalysis likely differ among DHFR enzymes.

DHFR is a model NADPH-dependent redox enzyme. Using conventional X-ray crystallography to study redox-sensitive enzymes is complicated by damage that results from the liberation of photoelectrons by incident X-rays. Prior work has established that proteins suffer site-specific X-ray-induced modifications at far lower absorbed doses than are required for more general radiation damage [30-34]. Metalloproteins are especially sensitive to X-ray photoreduction [35-38]. In some cases, this effect has been used to generate catalytically-relevant reducing equivalents that can drive metalloenzyme redox reactions in crystallo [39,40]. By contrast, there are few reports of non-metalloprotein enzyme chemistry being driven by X-rays in crystals, although functionally relevant radiation-driven changes to non-metalloprotein active sites have been observed [41-44].

In this study we analyze sub-1 Å resolution X-ray crystal structures of DHFR from *Bacillus subtilis* and *E. coli* bound to folate and NADP^+^. Electron density maps show evidence of changes in the active sites of both enzymes that are consistent with X-ray-driven hydride transfer. EcDHFR exhibits conformational heterogeneity that changes upon increasing radiation does and supports prior proposals for Met20-gated solvent access to the pterin ring of folate. BsDHFR has a different pattern of conformational heterogeneity, suggesting a distinct route to substrate protonation by solvent or possible crystal lattice artifacts. The broader implications of using X-ray-driven redox chemistry to study DHFR catalysis are discussed.

## Methods

### Protein purification and crystallization

EcDHFR was purified and crystallized as described previously [45]. The crystal used in this study was grown from the same batch of protein and the same NADP^+^ and folate stocks as the 100K and 277 K crystal structures reported in Keedy et al. (PDB 4PST, 4PSS, 4PTJ, 4PTH) [45] and Wan et al. (PDB 4PSY, 4RGC) [24]. BsDHFR was subcloned into pET22b from *B. subtilis* DNA templates that were previously constructed [46]. An insert bearing BsDHFR for subcloning was generated using PCR with the forward primer 5’-GTTGTTCATATGATTTCATTCATTTTTGCG-3’ and reverse primer 5’-GTTGTTACTAGTTTAAAATCCTCCCGCTTTAGA-3’. The amplification product was purified, digested with restriction enzymes, and ligated into pET22b. Native BsDHFR was purified by methotrexate affinity chromatography as reported previously [47]. BsDHFR in storage buffer (50 mM sodium phosphate pH=7.0, 10% glycerol, 2 mM DTT) was exchanged into 20 mM imidazole pH=7.0 and 1 mM folate (Sigma) by concentration and redilution and then concentrated to 25 mg/ml as determined using Bradford’s reagent (Bio-Rad). BsDHFR was supplemented with NADP^+^ (Sigma) to a final concentration of 6 mM, centrifuged to remove precipitate, and used for hanging drop sparse matrix crystallization screening at room temperature. Initial crystal hits were optimized at room temperature using hanging drop vapor diffusion and crystals were harvested from a condition with 10% PEG4000, 100 mM sodium acetate pH=5.0. Crystals were cryoprotected by passage through reservoir solution supplemented with PEG400 to a final concentration of 30% and cooled by immersion into liquid nitrogen.

### X-ray data collection, processing, refinement, and analysis

BsDHFR diffraction data were collected at the Stanford Synchrotron Radiation Laboratory (SSRL) beamline 11-1 at 100 K. Incident 0.9 Å X-rays illuminated a single crystal and diffraction data were collected on a ADSC Q315 charge-coupled device (CCD) detector. Different exposure times and crystal-to-detector distances were used to collect separate low- and high-resolution passes. Beamline geometry and incident X-ray energy constraints prevented collecting data beyond 0.93 Å resolution although the crystal diffracted beyond that limit judging from CC1/2 [48] and other data statistics (Table S1). Data were indexed and integrated in iMosfilm [49] and scaled in Aimless [50] in the CCP4 suite of programs [51]. Final data statistics are given in Table S1. The estimated absorbed radiation dose for this crystal is 9.9 MGy as calculated by RADDOSE [52].

EcDHFR diffraction data were collected at Advanced Photon Source (APS), BioCARS 14BM-C at 100 K. Incident X-rays at 0.9 Å illuminated a single crystal and diffraction data were collected on a ADSC Q315 charge-coupled device (CCD) detector. Different exposure times, crystal-to-detector distances, and detector offsets were used to collect separate low- and high-resolution passes within the dynamic range of the detector. As with BsDHFR, beamline geometry and incident X-ray energy constraints prevented collecting data beyond 0.93 Å resolution although the crystal diffracted beyond that limit. Data were indexed, integrated, and scaled in HKL2000 [53]. Final data statistics are given in Table S1. The estimated absorbed radiation dose for this crystal is 17.2 MGy as calculated by RADDOSE [52].

BsDHFR was phased by molecular replacement using EcDHFR (PDB 1RX2)[4] as a search model in CNS [54]. The model was manually improved in Coot [55], including building the ordered solvent model. Both EcDHFR and BsDHFR were refined against an intensity-based maximum likelihood target in PHENIX [56]. Anisotropic ADPs were refined for all non-hydrogen atoms and weights for stereochemistry and ADPs were optimized. Isomorphous difference maps between the 0.93 Å resolution 17.2 MGy EcDHFR dataset and the previously reported 0.85 Å resolution 4.9 MGy dataset (PDB 4PSY) [24] were calculated in PHENIX with no additional weighting. Final models were validated using tools in Coot [55] and MolProbity [57] and statistics are reported in Table S1. Anisotropic ADPs were analyzed using Rosenfield’s rigid body test as implemented in ANISOANL, part of the CCP4 suite of programs [51]. Differences of the projections of the anisotropic ADPs ellipsoids of pairs of atoms onto the line joining them were calculated for mainchain atoms and divided into 30 bins and averaged, corresponding to approximately five residues per bin. Structural figures were made with POVscript+ [58]. Multiple alignments were performed using Clustal Omega [59] and visualized in Jalview 2.11.2.7 [60].

## Results

### BsDHFR has a disulfide bond and a less dynamic active site loop than EcDHFR

The 0.93 Å resolution crystal structure of the BsDHFR-NADP^+^-folate pseudo-Michaelis complex shows that the enzyme is structurally similar to many other DHFRs. A DALI search [61] against the entire PDB identifies 760 PDB entries with an RMSD of 2.0 Å or better, all with Z-scores of 19 of higher. Many of these are multiple forms of the same enzymes, with *E. coli* (44% identity) and *S. aureus* (43% identity) DHFR being the highest-scoring DALI hits. Interestingly, BsDHFR is slightly more structurally similar to DHFRs from *S. aureus* (1.1 Å Cα RMSD; Z-score 27.4) [62] and *E. coli* (1.4 Å Cα RMSD; Z-score 27.4) [63] than it is to *B. anthracis* DHFR (1.4 Å Cα RMSD; Z-score 26.4) [28], despite the greater sequence identity between the two *Bacillus* enzymes (56% sequence identity).

We focus on a structural comparison with EcDHFR, as this is the most extensively studied bacterial DHFR. BsDHFR superimposes with EcDHFR over the entire protein length with a 1.4 Å Cα RMSD (Fig. 2A). Unlike EcDHFR, BsDHFR has a disulfide at C86-C92 linking an α-helix and β-strand, which is the most structurally divergent region of these enzymes (Fig. 2B). No other structurally characterized DHFR has a naturally occurring disulfide bond, although disulfides have been engineered into EcDHFR [64] and R67 DHFR [65]. Disulfides are uncommon in cytoplasmic proteins owing to the reducing environment of the cytosol, but they have been documented [66,67]. Conformational heterogeneity at the Cys86-Cys92 disulfide in BsDHFR indicates probable modification of this bond by partial X-ray photoreduction (Fig. 2C), as has been observed in other systems [30,43,44]. Cys86 and Cys92 are both present in DHFR sequences from close relatives of *B. subtilis* but are not well-conserved in sequences from other Bacillus species (Fig. 2D), indicating that the disulfide bond is not shared in most DHFRs in the genus Bacillus. Although the functional significance of the BsDHFR disulfide bond is unclear, it is remarkable that it persists during protein overexpression in an *E. coli* host possessing a highly reducing cytoplasmic environment.

**Figure 2.**
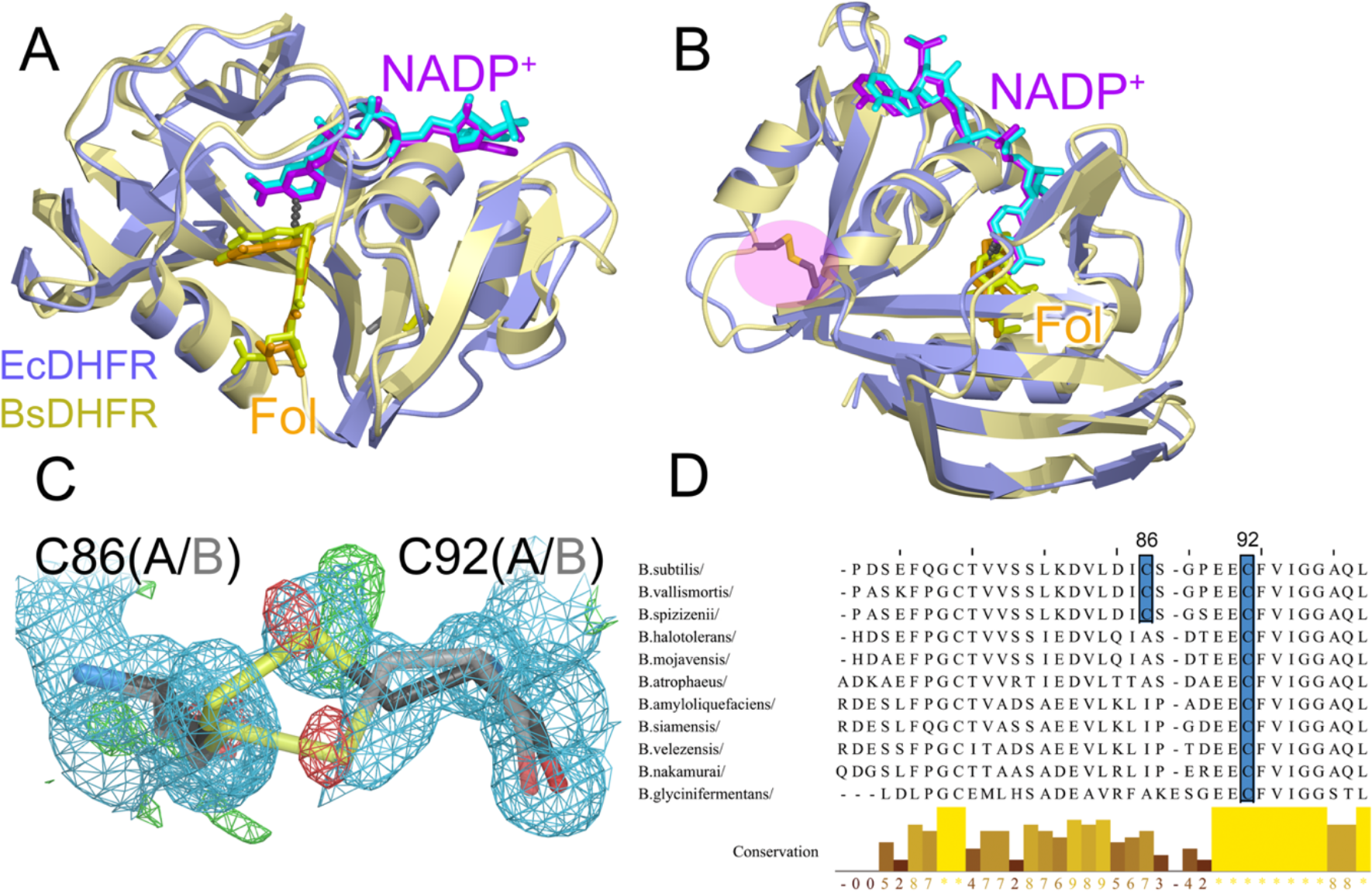
Structural comparison of EcDHFR an BsDHFR and presence of poorly conserved disulfide bond in BsDHFR. (A) Ribbon diagrams of the crystal structures EcDHFR (blue) and BsDHFR (yellow) are superimposed with a Ca RMSD of 1.4 Å. NADP^+^ and folate (Fol) are labeled, with purple NADP+ and orange Fol in EcDHFR and cyan NADP^+^ and yellow Fol in BsDHFR. (B) Another view of EcDHFR and BsDHFR superimposed, with the Cys86-Cys92 disulfide in BsDHFR shown in sticks and highlighted in pink. (C) The Cys86-Cys92 disulfide in BsDHFR with 2mFo-DFc electron density (blue) contoured at 0.8 α and mFo-DFc difference electron density contoured at +3α (green) and -3α (red). The disulfide is modeled in two conformations with the minor conformations in lighter sticks. The mFo-DFc peaks and conformational heterogeneity are evidence of partial photoreduction. (D) A sequence alignment of some Bacilli species, showing poor conservation of both cysteines (highlighted in blue and labeled) involved in the disulfide in BsDHFR (top sequence).

A notable difference between the active site regions of these enzymes is that Met20 in EcDHFR is replaced with Leu20 in BsDHFR. Conformational dynamics of the Met20 loop play an essential role in regulating NADP(H) cofactor binding and release during the EcDHFR catalytic cycle [4] and the EcDHFR Met20 loop shows evidence of disorder even in the closed conformation present in the NADP^+^-folate pseudo-Michaelis complex [23,24]. Motion of the Met20 sidechain is believed to be important for gating access of solvent to the reaction center during catalysis [8,23,24]. By contrast, the Leu20 loop in BsDHFR is well-ordered and shows no clear evidence of conformational heterogeneity. Visual inspection of Cα anisotropic atomic displacement parameters (ADPs) shows that the two enzymes have very different distributions of mainchain ADPs, which is supported by Rosenfield analysis of mainchain anisotropic ADPs (Fig. 3). Rosenfield analysis determines the difference in the projections of anisotropic ADPs of two atoms onto the line joining them. Atoms composing a rigid body should have identical anisotropic ADP projections onto the adjoining line, and thus the difference of their projection on this line is zero. Elevated values of this difference ADP projection indicate non-rigid body motion, consistent with greater internal flexibility. The EcDHFR Rosenfield matrix shows that the Met20 loop does not move as a rigid-body with the remainder of the EcHDFR (Fig. 3C), supporting prior work [4,8,23,24,26,45]. In contrast, anisotropic ADPs in the Leu20 loop of BsDHFR appear to be more correlated with ADPs in the rest of the protein, with lower difference ADP projection values in the Rosenfield matrix (Fig. 3D). BsDHFR crystallized at pH=5.0, and prior work noted that lower pH values increase the degree of order in the Met20 loop in EcDHFR [25]. Therefore, we cannot exclude the possibility that the Leu20 loop in BsDHFR may be more disordered at other pH values. Furthermore, there are lattice contacts near the BsDHFR Leu20 loop that are not present in EcDHFR, possibly restricting its conformational freedom.

**Figure 3.**
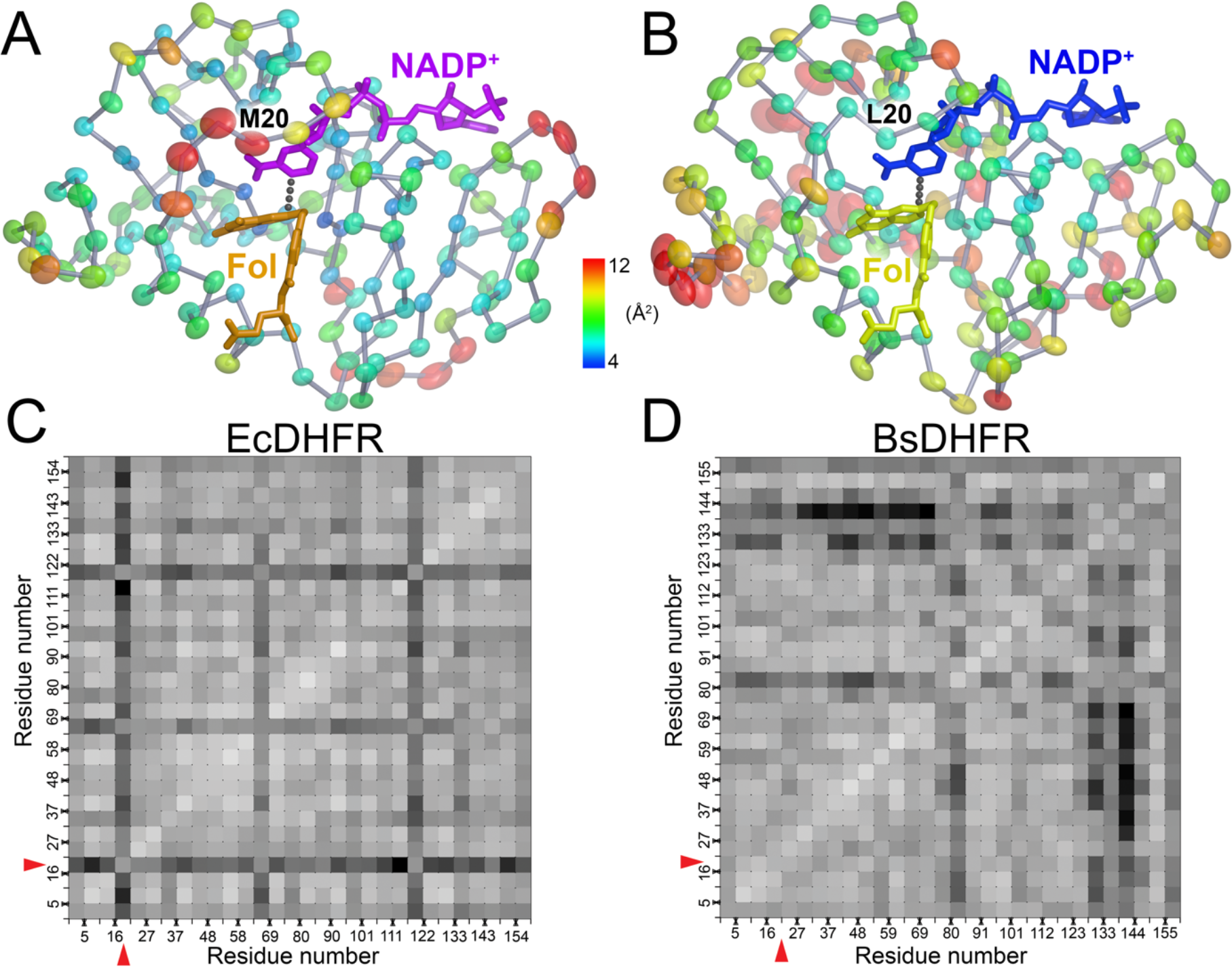
Anisotropic ADPs show differences in active site loop mobilities in EcDHFR and BsDHFR. (A) Ellipsoids for Cα atoms in EcDHFR are shown at the 95% probability level and colored from blue (4 Å^2^) to red (12 Å^2^). Met20 in the mobile active site loop is labeled. (B) Ellipsoids for Cα atoms in BsDHFR are shown at the 95% probability level and colored from blue (4 Å^2^) to red (12 Å^2^). Leu20 in the active site loop is labeled. (C) Rosenfield difference matrix for mainchain ADPs in EcDHFR shows non-rigid body motion (indicated by darker shades of grey) of the M20 loop (red arrow). (D) Rosenfield difference matrix for mainchain ADPs in BsDHFR shows largely rigid body character to the ADPS in the L20 loop (red arrow).

### X-ray photoreduction drives chemistry at the active site of DHFR crystals

A curious feature of the mFo-DFc electron density maps for BsDHFR is a positive peak spanning the C4 atom of NADP^+^ and the C6 atom of folate (Fig. 4A). These are the two atoms that would be involved in hydride transfer in the productive NADPH-dihydrofolate Michaelis complex and are only 3.16 Å apart in the BsDHFR-NADP^+^-folate crystal structure. However, the pseudo-Michaelis complex used here contains oxidized substrate and cofactor, so no chemistry should be possible in the crystal. Anisotropic ADPs in this region indicate motion of the nicotinamide and the pterin rings toward each other, roughly aligned with the extended mFo-DFc electron density feature (Fig. 4B). We did not observe similar difference density in the comparably high-resolution structures of EcDHFR at 100 K (0.85 Å, PDB 4PSY) or 277K (1.05 Å, PDB 4RGC). The BsDHFR crystal received an estimated absorbed dose of 9.9 megaGray (MGy), roughly twice the 4.9 MGy absorbed dose of the 4PSY EcDHFR crystal [24]. Therefore, we considered the possibility that this feature may be due to X-ray driven photoreduction of the previously oxidized abortive ternary complex.

**Figure 4.**
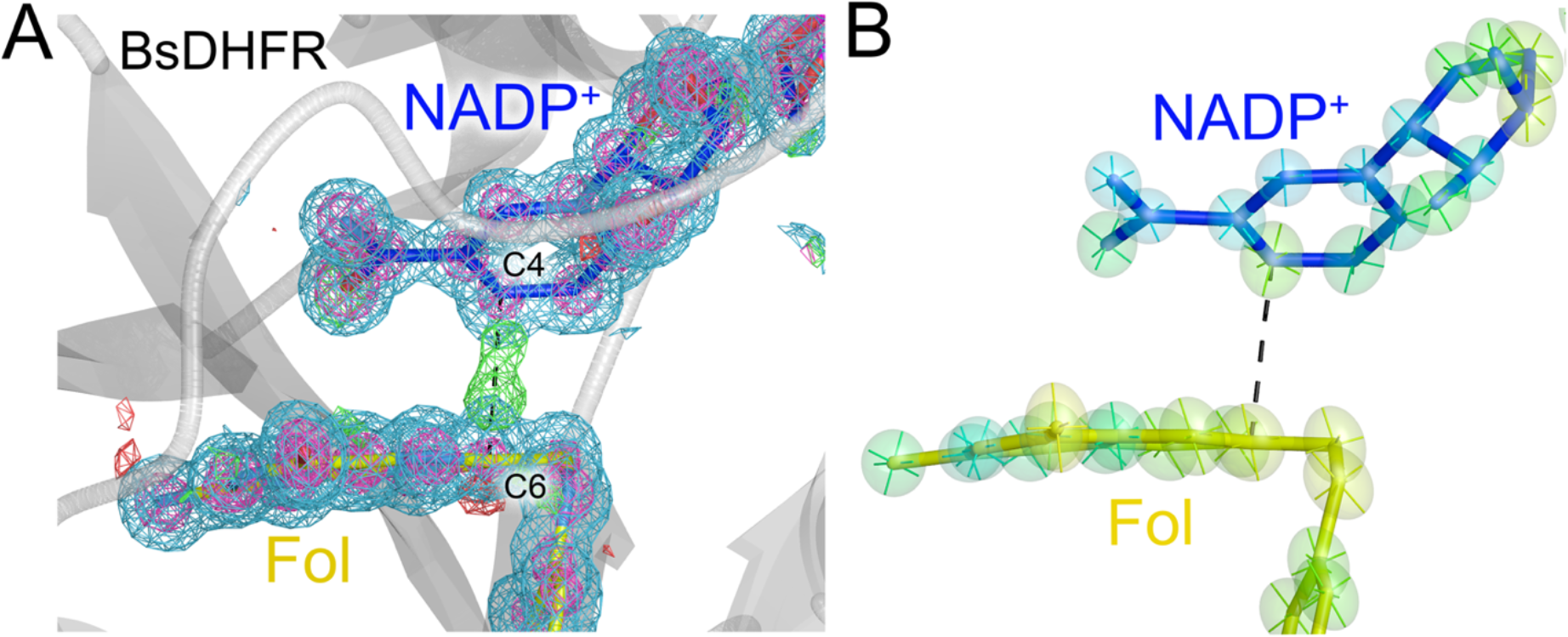
Electron density maps show evidence of hydride transfer in BsDHFR. (A) The BsDHFR active site with NADP^+^ (blue) and folate (yellow) is shown with 2mFo-DFc electron density contoured at 0.9α (blue) and 4.0α (purple) and mFo-DFc electron density contoured at +2.8α (green) and -2.8α (red). There is a strong positive peak in the mFo-DFc electron density map connecting the C4 atom of NADP^+^ and the C6 atom of folate, where hydride transfer would occur in the true Michaelis complex. (B) Anisotropic ADPs at the 60% probability level are colored from blue (5 Å^2^) to red (12 Å^2^) with principal axes shown. The anisotropic ADPs indicate displacements of the ring systems of NADP^+^ and folate that is roughly aligned with the positive mFo-DFc electron density feature in (A).

To test if X-ray photoreduction was driving chemistry in DHFR-NADP^+^-folate crystals, we turned to a previously unpublished 0.93 Å resolution structure of EcDHFR-NADP^+^-folate at 100 K. Beamline limitations prevented us from collecting data to the diffraction limit of the crystal, and data statistics indicate that it diffracts well beyond 0.93 Å resolution (Table S1). We collected this 0.93 Å resolution dataset prior to measuring a higher resolution 0.85 Å dataset at 100 K (PDB 4PSY) using a different data collection strategy that reduced the absorbed X-ray dose [24]. The crystal that produced the 0.93 Å dataset absorbed an estimated X-ray dose of 17.2 MGy as calculated in RADDOSE [52], over three times the absorbed dose of the 4PSY crystal and approximately 1.7x the dose of the BsDHFR dataset (see above). This crystal’s dose approaches but does not exceed the commonly quoted 30 MGy dose limit for cryocooled crystals [68]. The EcDHFR mFo-DFc electron density shows prominent evidence of changes in the folate molecule (Fig. 5A) that are similar to those observed in BsDHFR. An isomorphous difference electron density calculated between the higher- and lower-dose EcDHFR datasets (Fo(17MGy)-Fo(5MGy)) has peaks near the pterin ring that are correspond to the conformations of dideazaotetrahydrofolate (ddTHF) from PDB 1RC4 [4] and authentic tetrahydrofolate (PDB 6CW7) reported more recently [69] (Fig. 5B,C). Moreover, there are prominent Fo(17MGy)-Fo(5MGy) difference electron density peaks along the entire length of the NADP^+^ molecule (Fig. 5D) Because the 0.85 Å and 0.93 Å resolution EcDHFR datasets were collected from crystals grown from the same batch of enzyme, folate, and NADP^+^ under identical crystallization conditions, we conclude that X-ray driven chemistry is predominantly responsible for the partial formation of tetrahydrofolate (or a structurally similar species) in the active site of the 17.2 MGy EcDHFR dataset.

**Figure 5.**
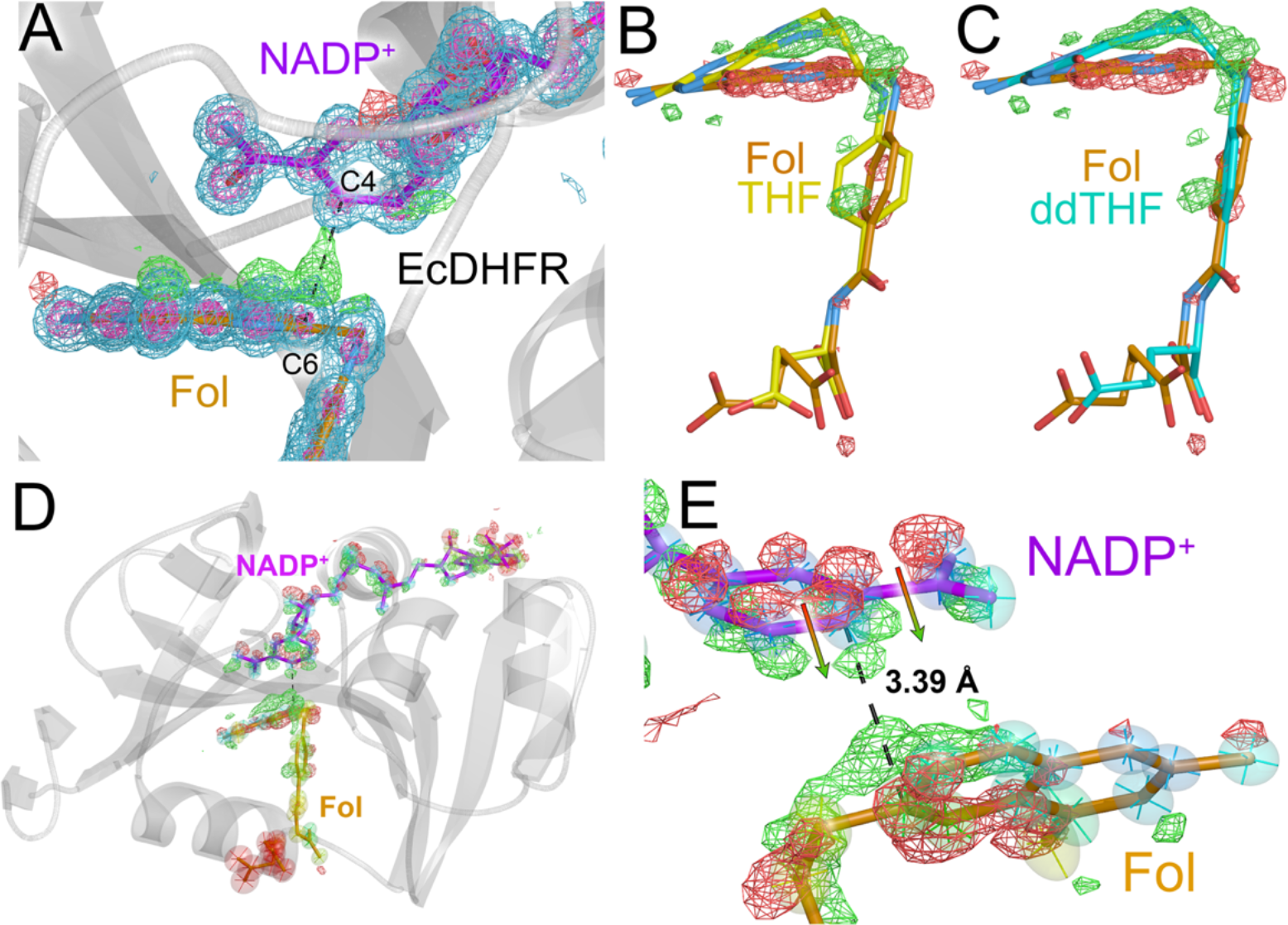
X-ray-driven hydride transfer and production of tetrahydrofolate in EcDHFR. (A) The EcDHFR active site with NADP^+^ (purple) and folate (orange) is shown with 2mFo-DFc maps contoured at 0.9σ (blue) and 4.0σ (purple) and mFo-DFc electron density contoured at +2.8σ (green) and -2.8σ (red). Several peaks in the mFo-DFc electron density map indicate modifications of the pterin that are similar to those seen in BsDHFR. (B) The Fo(17MGy)-Fo(5MGy) isomorphous difference map contoured at +3σ (green) and -3σ (red) shows partial conversion of folate to a species resembling tetrahydrofolate (THF). THF from PDB 6CW7 is shown in yellow sticks. (C) The Fo(17MGy)-Fo(5MGy) isomorphous difference map contoured at +3σ (green) and -3σ (red) shows conversion of folate to a species resembling the THF analog 5,10 dideazatetrahydrofolate (ddTHF) from PDB 1RC4 (cyan sticks). (D) The Fo(17MGy)-Fo(5MGy) isomorphous difference map (+3σ; green and -3σ; red) displayed over the entire NADP^+^ cofactor and folate molecules in EcDHFR, showing extended difference peaks over the whole NADP^+^ molecule. (E) The Fo(17MGy)-Fo(5MGy) isomorphous difference map (+3σ; green and -3σ; red) supports movement of the nicotinamide moiety of NADP^+^ towards pterin ring of folate upon increased X-ray dose. Arrows show direction of nicotinamide movement.

Atomically resolved Fo(17MGy)-Fo(5MGy) difference electron density features indicate an unexpected movement of the nicotinamide ring of NADP^+^ towards the partially reduced folate, and this movement is aligned with the principal axes of the anisotropic ADPs for these atoms (Fig. 5E). This is surprising, as formation of tetrahydrofolate should create greater steric conflict with the C4 atom of NADP^+^, and therefore would be expected to push the nicotinamide ring away from the pterin. Instead, the Fo(17MGy)-Fo(5MGy) electron density peaks indicate a compression of the distance between hydride donor and acceptor atoms. Considered in light of the close 3.39 Å contact between the hydride donor (NADP^+^ C4) and acceptor (folate C6) atoms in the active site, these observations indicate that chemistry closely resembling hydride transfer is occurring between these two atoms in the higher dose 17.2 MGy EcDHFR crystals and is likely also occurring in BsDHFR crystals.

The discovery of radiation-driven chemistry similar to that natively catalyzed by DHFR provides a means to observe conformational changes that may be occurring during EcDHFR catalysis. The Met20 sidechain is modeled in two conformations in this structure (Fig. 6A), similar to prior work [24] and further informed by a recent detailed study of EcDHFR [23]. In one conformation, the Cε methyl group occludes the active site, while the other conformation permits entry of a partially occupied water molecule as noted by Wan et al [24]. Consequently, the water molecule is modeled at 0.47 occupancy which closely matches the 0.48 occupancy of the permissive Met20 conformation. Fo(17MGy)-Fo(5MGy) electron density indicates a redistribution of the populations of these Met20 sidechain conformations upon increased radiation dose, with the occluded conformation becoming less populated and the conformer that permits water entry becoming more populated (Fig. 6B). This is the same motion that we and others have proposed assists dihydrofolate protonation during the EcDHFR catalytic cycle [20,23-25]. Notably, the partially occupied water molecule near Met20 in EcDHFR is not present in BsDHFR. Therefore, BsDHFR likely uses a distinct structural mechanism for substrate protonation.

**Figure 6.**
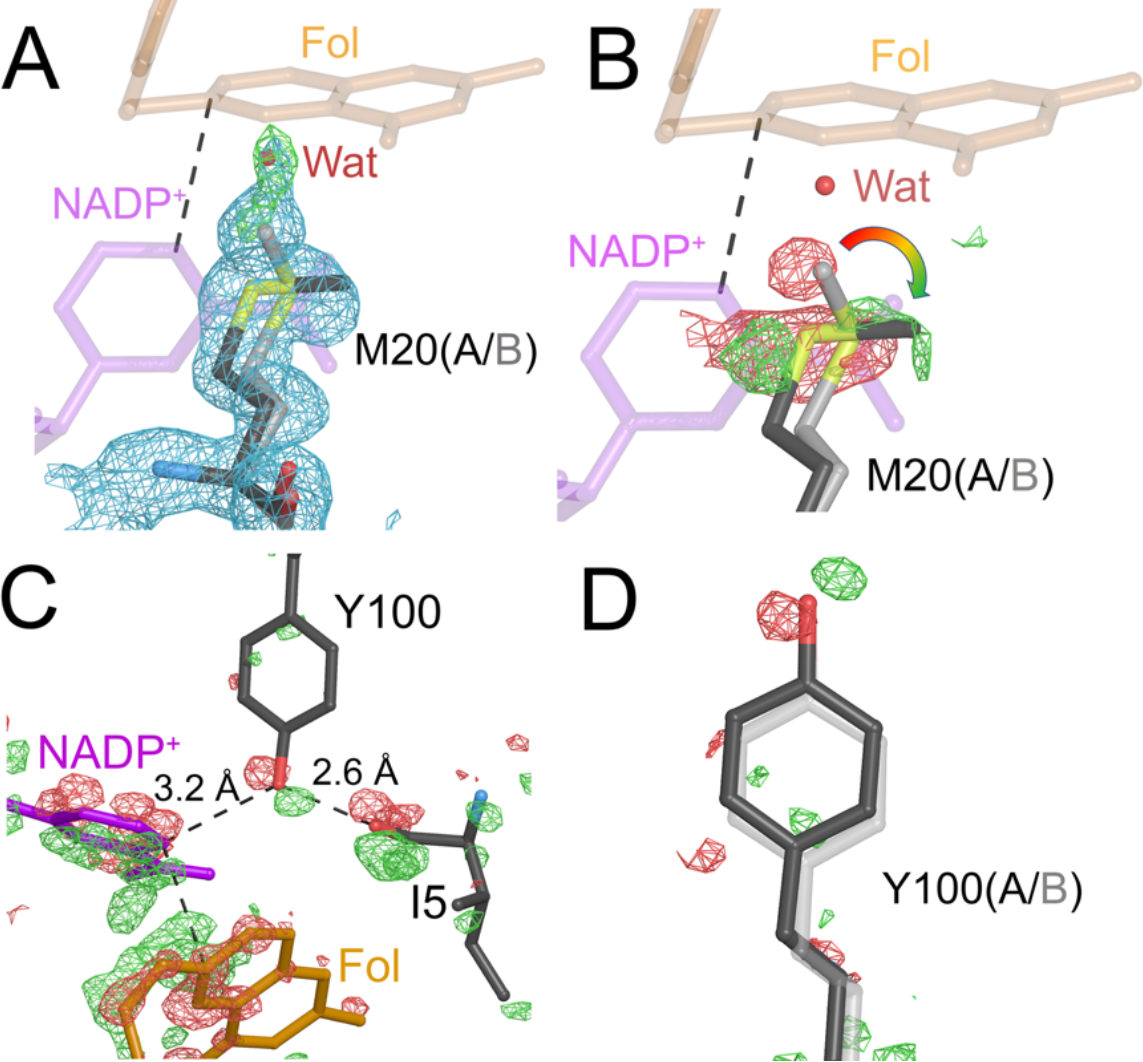
Increased X-ray dose results in Met20 motion consistent with solvent gating. (A) 2mFo-DFc electron density contoured at 0.7σ (blue) and polder omit mFo-DFc at contoured at 4.0σ (green) show a partially occupied water molecule (red, labeled) whose occupancy is correlated with conformational disorder at Met20. The water molecule can only occupy the site if Met20 is in the permissive (black) conformation. (B) The Fo(17MGy)-Fo(5MGy) isomorphous difference map (+3σ; green) and -3σ; red)) shows that Met20 shifts into the permissive conformation upon increased X-ray dose. (C) The Fo(17MGy)-Fo(5MGy) isomorphous difference map (+3σ; green, -3σ; red) shows that Tyr100 dose-dependent motion is spatially correlated with motion of the NADP^+^ nicotinamide and surrounding residues. (D) The EcDHFR Fo(17MGy)-Fo(5MGy) isomorphous difference map (+3σ; green and -3σ; red) has peaks for Tyr100 movement that roughly agree with modelled disorder at this residue in BsDHFR. The more occupied conformation is in the darker line.

Other residues in the active site also move in response to increased radiation dose. Tyr100 moves in the same direction as the nicotinamide ring of NADP^+^ (Fig. 6C) and nearby residues, including Ile5. Tyr100 has been suggested to be electrostatically and dynamically coupled to catalysis in EcDHFR [70] and is conserved in BsDHFR, where it is modeled in two conformations that are displaced in a similar direction as the Fo(17MGy)-Fo(5MGy) difference electron density for EcDHFR (Fig. 6D). Several peaks in this difference electron density map extend along the NADP^+^ molecule and radiate outwards to surrounding protein residues, suggesting that the entire NADP^+^ cofactor and its binding site is conformationally responsive to hydride transfer/photoreduction.

### Other sites of radiation-driven changes in EcDHFR

There are many strong (>4σ) peaks in the Fo(17MGy)-Fo(5MGy) electron density map, not all of which are easily interpreted as being the direct result of chemistry at the site of hydride transfer (Fig 7A). Cys152 is oxidized to a sulfonate (-SO3-) in the 0.93 Å resolution EcDHFR structure (Fig. 7B), a higher oxidation state than the sulfinate (SO2-) that was modeled in PDB 4PSY [24]. This more extensive cysteine oxidation, which increases the occupancies of all three oxygen atoms, results in distributed changes in the surrounding protein and nearby solvent. Certain cysteine residues are known to be sensitive to radiation-induced oxidation [71], and these data provide a detailed view of how photooxidative modification of a cysteine residue can result in correlated changes to its protein microenvironment over several Ångstroms. Other changes that are not obviously connected to photoreduction of the active site include peaks near the Mn^2+^ sites. The largest single peak (42.5 α) in the Fo(17MGy)-Fo(5MGy) difference electron density is on Mn^2+^203, which is coordinated by His149 on the same β-strand as Cys152. Although metals are preferred sites for radiation damage, it is not clear how increased radiation dose would cause an elevation in Mn^2+^ occupancy or a strong positive Fo(17MGy)-Fo(5MGy) difference peak, leaving open the possibly that this feature is due to sample variability between the two crystals used to calculate the isomorphous difference map. Despite the numerous and strong peaks in the Fo(17MGy)-Fo(5MGy) difference electron density map, the refined mainchain ADPs for this EcDHFR model are nearly identical to those of PDB 4PSY (Fig.7C). This close agreement in refined mainchain ADP values demonstrates that widespread general radiation damage is not occurring the 17 MGy crystal and is remarkable given that 4PSY refined with SHELXL [72] while the present 17 MGy model was refined with PHENIX [56], which use different types of ADP restraints and refinement target functions.

**Figure 7.**
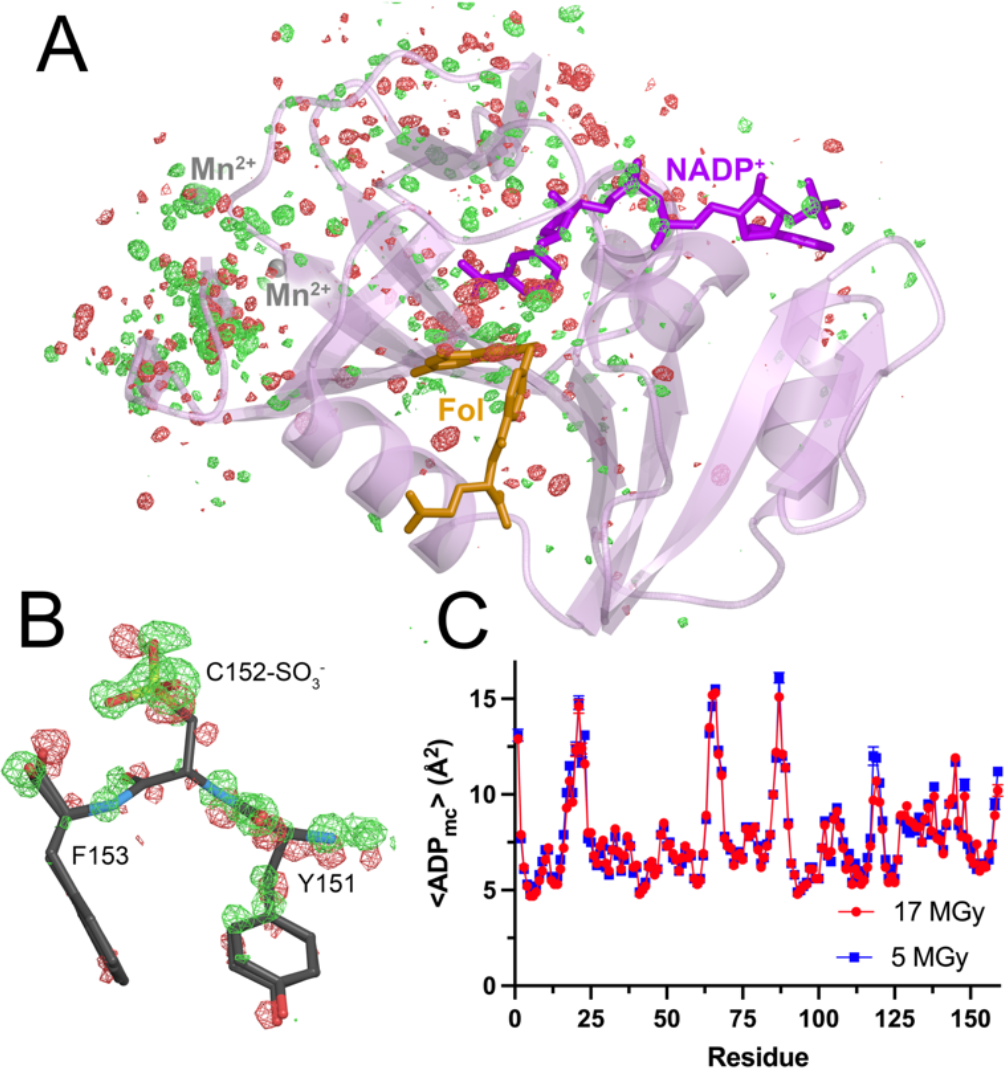
Radiation dose-dependent changes at other locations in EcDHFR. (A) The Fo(17MGy)-Fo(5MGy) isomorphous difference map (contoured at +4σ; green and - 4σ; red) for the entire EcDHRR-NADP^+^-folate complex shows many peaks that are distal to the active site. (B) The Fo(17MGy)-Fo(5MGy) isomorphous difference map (+3σ; green and -3σ; red) shows increased dose-dependent oxidation at Cys152 that causes movement in surrounding residues. (C) A plot of averaged mainchain ADPs for 17MGy and 5MGy EcDHFR models are nearly identical, indicating that there are no marked global changes in ADPs that would indicate global radiation damage.

## Discussion

X-rays profoundly alter the redox state of irradiated crystals and are well-known to cause chemical changes in proteins [34,36,38,73-76]. In some cases, these chemical changes resemble the reactions catalyzed by redox-active enzymes, particularly in metalloenzymes [39,40]. X-ray photoreduction of enzyme active sites can cause difficulties in interpretation and has been a motivation for collecting data using X-ray free electron lasers where femtosecond pulse of incident X-rays result in minimal radiation damage [77-79]. DHFR is a rare example of a non-metalloenzyme that displays X-ray-driven catalysis in the crystal. It seems likely that such reactions are more common than has been reported and that inspection of other non-metalloprotein redox enzyme crystal structures may uncover additional evidence of X-ray driven redox chemistry that recapitulates the native enzyme-catalyzed reaction. It is our hope that this report motivates additional scrutiny of other redox enzyme crystal structures as a function of absorbed dose. Although there are many caveats about the differences between X-ray photoelectron-driven chemistry and native enzyme catalysis, it is possible that in favorable cases X-ray-driven reactions can provide a window into catalysis-coupled protein conformational dynamics.

Features in the Fo(17MGy)-Fo(5MGy) EcDHFR difference electron density map support prior suggestions that the conformational dynamics of Met20 in the closed loop conformation are coupled to catalysis [23-25,80]. Our analysis does not indicate how such coupling occurs, although the significant change in pterin geometry that occurs upon tetrahydrofolate formation [81] or concomitant protonation of the N5 atom of the pterin are plausible hypotheses. In contrast, BsDHFR lacks Met20 and the loop containing Leu20 is not more dynamic than surrounding residues in this P61 crystal form. Like BsDHFR, human DHFR also has a more rigid active site loop with Leu22 instead of Met20 [26], and Greisman et al have suggested that dynamics at a different residue (Phe31) gates solvent access to the human DHFR active site [23]. BsDHFR has Leu28 instead of Phe31 and no obvious evidence of Leu28 mobility at 100K, suggesting that solvent-mediated substrate protonation occurs via a different route in BsDHFR or that the low pH or restrictive lattice contacts in this crystal form prohibit full manifestation of functionally relevant coupled protein-solvent dynamics. Additional studies, preferably with structures of BsDHFR crystallized in multiple space groups and measured at room temperature, will be needed to resolve this issue.

We observe dose-dependent conformational shifts in EcDHFR Tyr100 that are similar to those of the nicotinamide ring of NADP^+^, broadly consistent with prior proposals of a role for Tyr100 in stabilizing electrostatic changes in the EcDHFR active site during catalysis [16,70]. Unlike Met20, Tyr100 is conserved in BsDHFR. The area near Tyr100 (including Trp22 and Phe125) in the Fo(17MGy)-Fo(5MGy) electron density map has features that resemble difference electron density maps calculated between datasets collected at different temperatures in Greisman et al., indicating that distinct types of perturbation may cause similar correlated displacements in these residues [23]. The most unexpected change arising from greater absorbed X-ray dose in EcDHFR crystals was the movement of the NADP^+^ nicotinamide ring toward the pterin moiety of the folate. This movement has not been previously reported, although it must occur to facilitate hydride transfer in the true Michaelis complex of NADPH and dihydrofolate, where the transition state C4-C6 interatomic distance is predicted to be ~2.7 Å [18-20]. Compression of the nicotinamide C4-folate C6 interatomic distance supports the conclusion that catalytically relevant chemistry is occurring in the EcDHFR active site upon X-ray irradiation. Moreover, this result lays the groundwork for more detailed X-ray diffraction studies of bacterial DHFRs as a function of in multiple absorbed dose series. Direct comparison of dose-dependent datasets with existing temperature-dependent datasets [23] could be particularly informative.

BsDHFR contains a poorly conserved disulfide between C86 and C92, which is unexpected in a cytoplasmic protein. *B. subtilis* does not have an elevated number of proteins containing even numbers of cysteine residues [82], suggesting that cytoplasmic disulfide-containing proteins are rare in that organism. The disulfide is distal from the BsDHFR active site and does not have an obvious catalytic or structural role, raising questions about why it is present. *B. subtilis* and other Firmicutes use bacillithiol as the dominant intracellular small molecule thiol, which has a somewhat higher redox potential (−221 mV) than glutathione (−240 mV), the more common small molecule thiol present in *E. coli* and other organisms [83,84]. However, this disulfide formed when recombinant BsDHFR is expressed in an *E. coli* host, indicating that the details of the native intracellular small molecule thiol are not essential to its formation. Artifactual oxidation during protein purification is unlikely given the presence of 2 mM DTT in the storage buffer, and thus we propose that the BsDHFR disulfide likely has an atypically low redox potential whose structural determinants merit further investigation.

## Conclusions

X-rays generate photoelectrons that can facilitate a wide variety of chemical reactions inside crystals during X-ray diffraction data collection. Abortive ternary complexes of *B. subtilis* and *E. coli* DHFR bound to NADP^+^ and folate show evidence of hydride transfer that recapitulates the native reaction catalyzed by DHFR, NADPH, and dihydrofolate, including the production of tetrahydrolate (or a structurally similar species) in EcDHFR at higher X-ray dose. We conclude that X-rays can drive catalytically relevant chemistry in DHFR and possibly in other redox enzymes that do not contain metals. The changes that we observe in EcDHFR conformational heterogeneity upon increased X-ray dose are similar to previous reports of EcDHFR dynamics using diverse methods, suggesting that dose-dependent studies of redox enzymes may provide a useful way of observing conformational dynamics that occur during catalysis. However, electrons generated by X-rays behave differently than those delivered to redox enzymes by physiologically relevant reductants and thus conclusions drawn from X-ray-driven reactions should be viewed with caution until confirmed using complementary approaches. Our comparisons of BsDHFR and EcDHFR revealed conserved aspects of catalysis as well as uncovering idiosyncratic features of this important class of enzymes, including an unexpected disulfide in BsDHFR. The structural biology of bacterial DHFRs continues to surprise even after decades of study.

## Dedication

This article is dedicated to the memory of Prof. Chris Dealwis (Case Western Reserve University), whose many contributions to structural biology included important work on dihydrofolate reductases. M.A.W. was fortunate to collaborate with Chris and his group on some studies of *E. coli* DHFR.

## Author contributions

N.S. analyzed data, A.R.H. performed experiments, M.A.W. performed experiments, analyzed data, and wrote the manuscript.

## Data Availability Statement

Structure factors and refined model coordinates have been deposited with the PDB with accession codes 8UVZ (BsDHFR-NADP^+^-folate) and 8UW0 (EcDHFR-NADP^+^-folate).

### Acknowledgements

We thank Steven Benkovic (Pennsylvania State University) and Greg Petsko (Brandeis University) for early support of this work and Vukica Srajer and Robert Henning (Advanced Photon Source) for useful information about beamline 14BM-C’s configuration.

