## Supplemental Table 1 for "X-ray-driven chemistry and conformational heterogeneity in atomic resolution crystal structures of bacterial dihydrofolate reductases"

**Table S1: Crystallographic data and model statistics**

| Sample | EcDHFR-NADP$^+$-folate | BsDHFR-NADP$^+$-folate |
|---|---|---|
| Diffraction source | APS 14BM-C | SSRL 11-1 |
| Wavelength (Å) | 0.900 | 0.900 |
| Temperature (K) | 100 | 100 |
| Detector | ADSC Q315 | ADSC Q315 |
| Space group | $P2_12_12_1$ | $P6_1$ |
| a, b, c (Å), α, β, γ (°) | 33.98, 44.85, 98.24, 90, 90, 90 | 88.45, 88.45, 37.27, 90, 90, 120 |
| Mosaicity (°) | 0.24 | 0.26 |
| Resolution range (Å) | 33.12-0.93 (0.96-0.93) | 44.22-0.93 (0.95-0.93) |
| Total no. of observations | 525029 (24006) | 1245117 (50266) |
| No. of unique observations | 99062 (9233) | 110282 (5386) |
| Completeness (%) | 97.2 (91.7) | 100 (99.1) |
| Multiplicity | 5.3 (2.6) | 11.3 (9.3) |
| ⟨I/σ(I)⟩ | 29.6 (8.1) | 20.9 (2.8) |
| $CC_{1/2}$[1] | 0.999 (0.958) | 0.999 (0.873) |
| $R_{meas}$[3] | 0.050 (0.208) | 0.065 (0.657) |
| Model statistics | | |
| PDB code | 8UW0 | 8UVZ |
| Refinement program | PHENIX 1.19.2_4158 | PHENIX 1.19.2_4158 |
| Resolution | 33.12-0.93 (0.94-0.93) | 44.22-0.93 (0.94-0.93) |
| No. of reflections | 99005 (3938) | 110248 (3415) |
| No. of reflections, test set | 3003 (128) | 5504 (177) |
| $R_{work}$ | 0.1004 (0.1182) | 0.1103 (0.2015) |
| $R_{free}$ | 0.1152 (0.1317) | 0.1233 (0.2283) |
| No. of non-H atoms | | |
| Protein | 1796 | 1814 |
| Water | 371 | 301 |
| Heteroatoms | 88 | 110 |
| Total | 2255 | 2225 |
| Average R.M.S. deviations | | |
| Bonds (Å) | 0.012 | 0.013 |
| Angles (°) | 1.564 | 1.671 |
| Average B factors (<$B_{iso}$>;Å$^2$) | | |
| Protein | 8.98 | 11.12 |
| Water | 23.39 | 24.55 |
| Heteroatoms | 9.13 | 10.82 |
| Average ADP anisotropy[1] | | |
| Protein | 0.513 | 0.511 |
| Water | 0.383 | 0.510 |
| Heteroatoms | 0.528 | 0.585 |
| MolProbity clashscore | 2.5 | 2.2 |
| Ramachandran plot | | |
| Outliers; Allowed; Favored (%) | 0; 1.95; 98.05 | 0; 0.63; 99.37 |
